# An easily modifiable conjugative plasmid for studying horizontal gene transfer

**DOI:** 10.1101/2022.03.09.483620

**Authors:** Qinqin Wang, Asmus Kalckar Olesen, Lorrie Maccario, Søren J. Sørensen, Jonas Stenløkke Madsen

**Affiliations:** Section of Microbiology, Department of Biology, University of Copenhagen, 2100 Copenhagen, Denmark

## Abstract

Horizontal gene transfer is an important mechanism in bacterial evolution and can occur at striking frequencies when mediated by mobile genetic elements. Conjugative plasmids are mobile genetic elements that are main drivers of horizontal transfer and a major facilitator in the spread of antibiotic resistance genes. However, conjugative plasmid models that readily can be genetically modified with the aim to study horizontal transfer are not currently available. The aim of this study was to develop a conjugative plasmid model where the insertion of gene cassettes such as reporter genes (*e.g*., fluorescent proteins) or antibiotic resistance genes would be efficient and convenient. For that, we introduced a single *att*Tn7 site into the conjugative broad-host-range IncP-1 plasmid pKJK5 in a non-disruptive manner. Furthermore, a version with lower transfer rate and a non-conjugative version of pKJK5-*att*Tn7 were also constructed. The advantage of having the *att*Tn7 sites is that genes of interest can be introduced in a single step with very high success rate using the Tn7 transposition system. In addition, larger genetic fragments can be inserted. To illustrate the efficacy of the constructed pKJK5 plasmids, they were complimented with *sfgfp* in addition to seven different β-lactamase genes representing the four known classes of β-lactamases.

## Introduction

Horizontal gene transfer (HGT) is an important facilitator of bacterial evolution and when mediated by mobile genetic elements, such as conjugative plasmids, HGT can occur at striking frequencies (Sørensen et al., 2005). Conjugative plasmids are especially known for their role in spreading antibiotic resistance genes (Che et al., 2021) and virulence factors (Ghigo, 2001), however, while this underlines their importance, there are still many unknowns about the biology of plasmids and HGT in general. Here we constructed a conjugative plasmid which easily can be complimented with genetic cassettes to investigate various aspects of HGT including plasmid biology, transfer dynamics, accessory genes, stability, etc. Molecular engineering of conjugative plasmids can have numerous objectives such as, complimenting conjugative plasmids with reporter genes that can be used to study transfer dynamics at the single cell level (Pinilla-Redondo et al., 2018), testing what genes can be transferred stably by conjugation, factors that affect the success of HGT, and metagenome engineering and cargo delivery by conjugation. Though, many other applications can be imagined.

Tn7 transposons have been widely used in bacterial genome engineering enabling gene complementation and expression (Peters and Craig, 2001). When transposition is mediated by TnsABC+D the Tn7 transposon can be inserted specifically into chromosomes at an *att*Tn7 site, which typically is located downstream of the bacterial glutamine synthetase (*glmS*) gene of gram-negative bacteria which is highly conserved (Peters and Craig, 2001). Choi et al. constructed the widely used mini-Tn7 system where the genetic material one wishes to transpose is flanked by inverted repeats named Tn7L and Tn7R on a transfer plasmid. Tn7L and Tn7R are recognized by the transposase complex TnsABCD, encoded on the helper plasmid, and then inserted at the chromosomal *att*Tn7 (Choi et al., 2005). The bacterial Tn7-transposon-based delivery systems have important advantages: (i) It has high affinity integration at *att*Tn7 sites; (ii) both small and large DNA fragments can be integrated; (iii) integration at a defined target site can be done without any deleterious effects or preparatory genetic modification; and (iv) the inserted transposon is maintained without antibiotic selection (Choi et al., 2005; Choi and Schweizer, 2006; Choi and Kim, 2009; Remus-Emsermann et al., 2016). Later derivatives of the mini-Tn7 system include pGRG36 where Tn7L and Tn7R associated multiple cloning sites and the transposase complex TnsABC+D were integrated into a single temperature-sensitive vector, which furthered the ease by which Tn7-based gene complementation can be done (McKenzie and Craig, 2006).

Another efficient gene integration system is the λ phage derived Red recombination system, a mutagenesis method that through homologs recombination allows defined insertions of genes, deletions, or point mutations in bacteria and fungi (Chaveroche et al., 2000; Datsenko and Wanner, 2000). With this method, efficient recombination can be achieved between polymerase chain reaction (PCR) products and a target replicon by induction of the λ phage Red operon, provided that the linear DNA obtained by PCR has ca. 36-nt or larger flanking extensions that are homologous to the target DNA. The Red operon encodes three genes, *gam, exo* and *beta*. Gam prevents the intracellular exonucleases from digesting the linear DNA introduced into the bacteria. Exo will degrade linear dsDNA starting from the 5’ end and generate ssDNA, and Beta protects the ssDNA produced by the Exo and promotes its annealing to the complementary ssDNA target in the replicon (Datsenko and Wanner, 2000). The great advantage of the Red recombination system is that mutagenesis can be done specifically at any genomic site (Datsenko and Wanner, 2000).

Here we utilized the λ Red recombination system to inset an *att*Tn7 site into a model conjugative plasmid pKJK5, in a non-disruptive manner. pKJK5 belongs to the IncP-ε incompatibility group (Bahl et al., 2007a) and has a very broad-host ranges (Klümper et al., 2015). It transfers at high frequencies to many proteobacteria but has also been reported to transfer to gram-positive bacteria (Klümper et al., 2015) and IncP plasmids have been demonstrated to transfer even to eukaryotes (Hayman and Bolen, 1993). After successful insertion of *att*Tn7 into pKJK5, generating pKJK5-*att*Tn7, we demonstrate the ease at which genes can be complemented into pKJK5-*att*Tn7 where only a few PCR and screening steps are needed to achieve the desired results. In brief, the model conjugative plasmid presented here can be engineered easily and reliably to advance research on HGT and can in general help advance studies by efficiently decreasing time spent engineering and reducing the reagents and equipment needed.

## Methods

### Plasmids and strains

The bacterial strains and plasmids used in this study are listed in **Table 1**. *Escherichia coli* strains were grown in Luria-Bertani (LB) medium at 30°C or 37°C (Green and Sambrook, 2012). Antibiotics (Sigma-Aldrich) were used at the following concentration: Ampicillin (Amp, 100 μg/ ml), kanamycin (Kan, 100 μg/ml), tetracycline (Tet, 15 μg/ml), gentamicin (Gen, 10 μg/ml), rifampicin (Rif, 100 μg/ ml), nalidixic acid (Nal, 100 μg/ ml), chloramphenicol (Chl, 30 μg/ml), Cefotaxime (Ctx, 2 μg/ml), and Meropenem, (Mem, 0.5 μg/ml)). Plasmid DNA was extracted from overnight cultures using the Plasmid Mini AX kit (A&A Biotechnology). Primers (TAG Copenhagen A/S) used in this study are in **Table 2**.

**Table 1.**
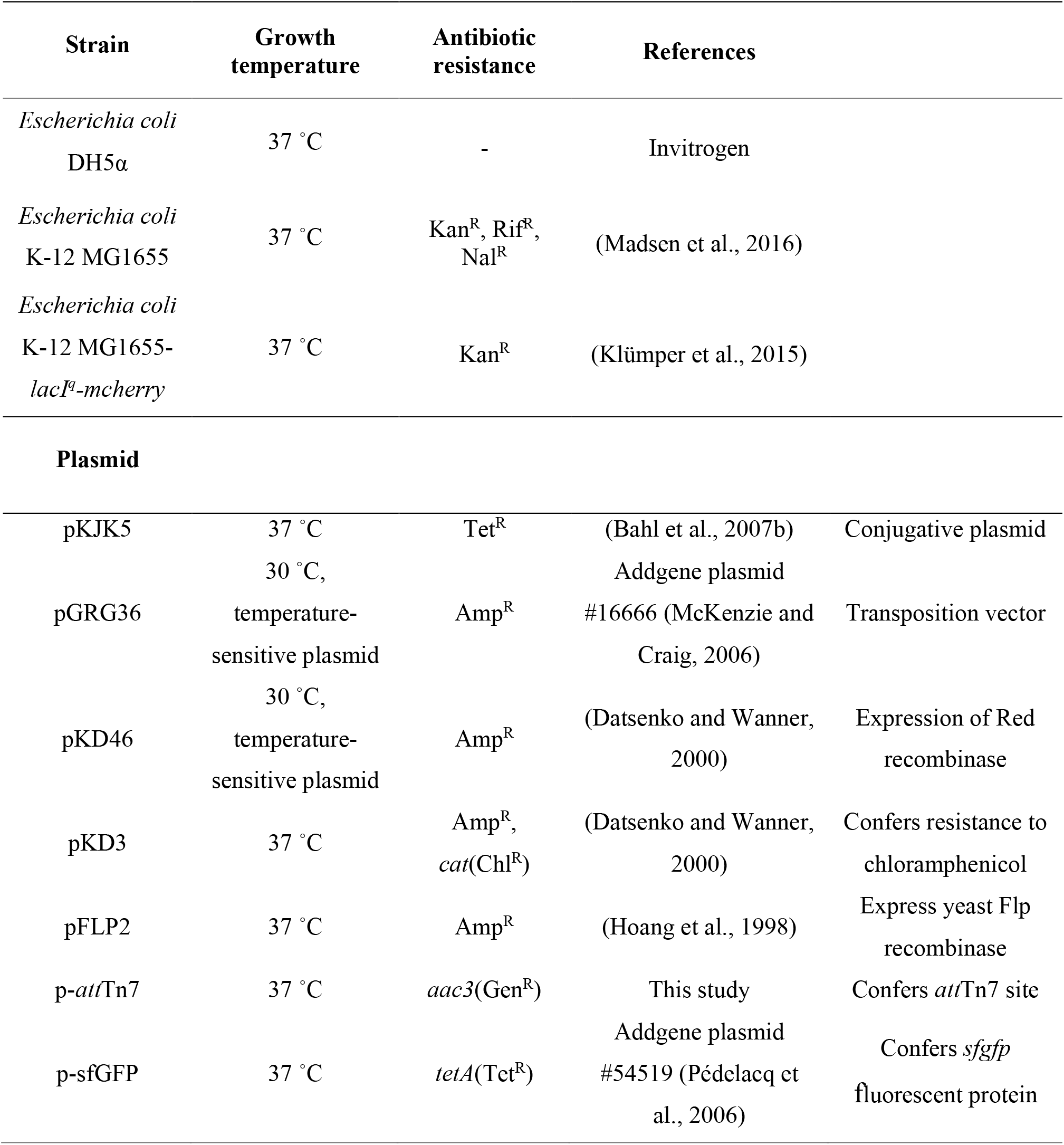
Strains and plasmids used in this study.

**Table 2.**
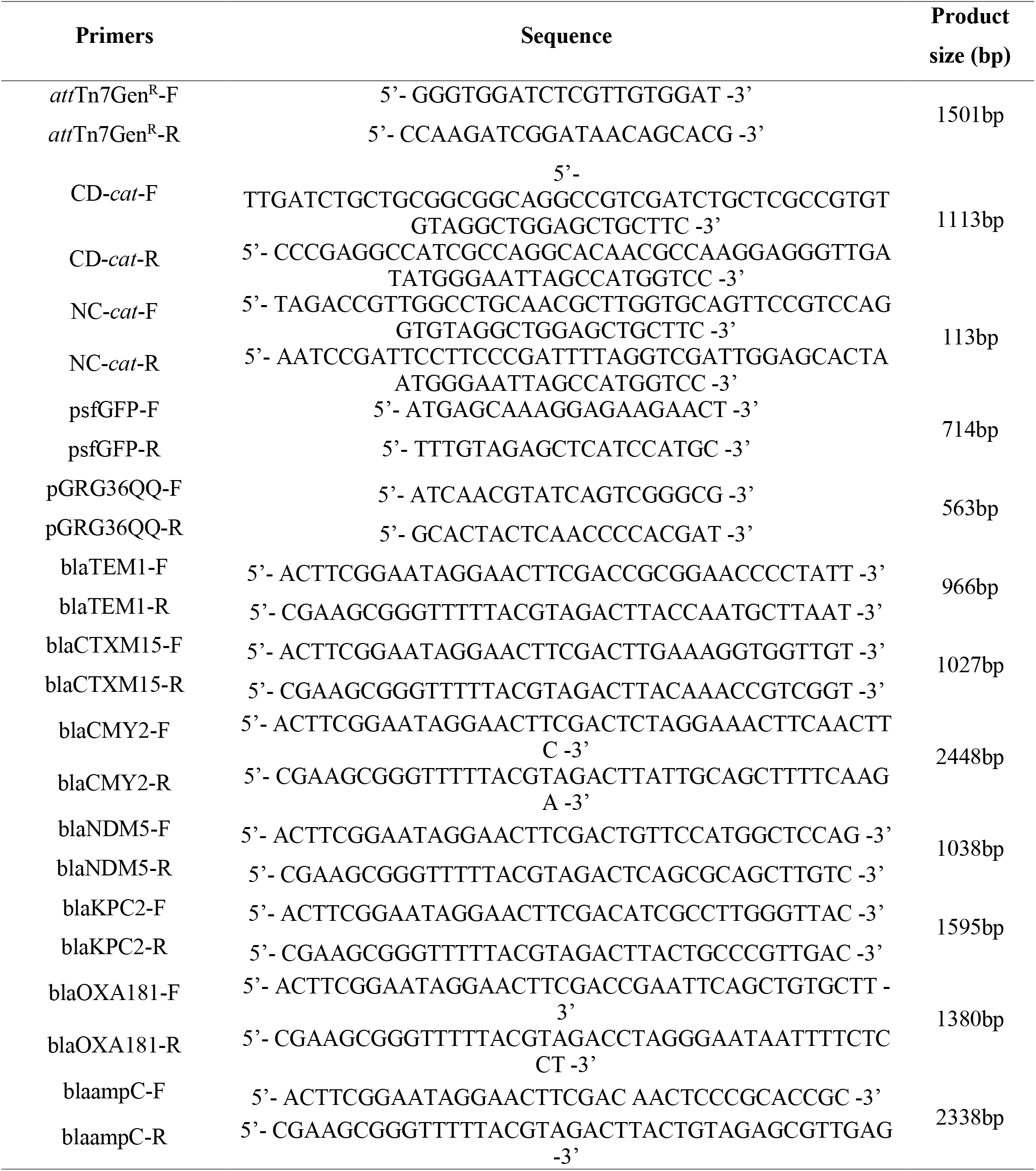
Oligonucleotide primers used in this study.

### Construction of plasmid pKJK5-*att*Tn7

p-*att*Tn7 was synthesized by ThermoFisher Scientific based on a provided design, as further described in the results section (**Fig. 1**). From p-*att*Tn7 the *att*Tn7-*aac3* fragment, including *FRT*-sites, was amplified by PCR using primers *att*Tn7Gen^R^-F and *att*Tn7Gen^R^-R that included 40bp overhangs with homology to the pKJK5 target region. Next, λ Red recombination was used (Serra-Moreno et al., 2006). 100-150 ng of the linear PCR product was transformed by electroporation (1.8 kV for ∼5 ms) into competent *E. coli* DH5α cells already harboring both the helper plasmid pKD46 and the target plasmid pKJK5. The selection was done on LB agar medium containing Amp, Tet, and Gen. Growing colonies were screened by colony PCR using the *att*Tn7Gen^R^-F and *att*Tn7Gen^R^-F primers. Positive colonies were grown in LB with Tet and Gen at 42°C overnight to remove pKD46. Post plasmid curing, it was ensured that colonies were sensitive to Amp. Hereafter, the helper plasmid pFLP2 was transformed into the *E. coli* DH5α cells containing the now constructed pKJK5-*att*Tn7*-aac3* by electroporation, to obtain plasmid pKJK5-*att*Tn7 (Choi and Schweizer, 2005). Clones that grew on LB agar medium with Tet but didn’t grow in the presence of Gen were selected and verified by colony PCR with primers *att*Tn7Gen^R^-F and *att*Tn7Gen^R^-F. After the flipase mediated removal of *aac3*, we used LB agar plates containing 5 g/ml sucrose to select for clones without pFLP2, and solely pKJK5-*attTn7*. pKJK5-*att*Tn7 was whole-genome sequenced by Nanopore sequencing carried out on R9.4 MinION flowcells (Oxford Nanopore Technologies) for up to 48 h. Libraries were prepared using the Rapid Barcoding Sequencing Kit (Oxford Nanopore Technologies, SQK-RBK004) following the manufacturer’s instructions.

**Fig. 1.**
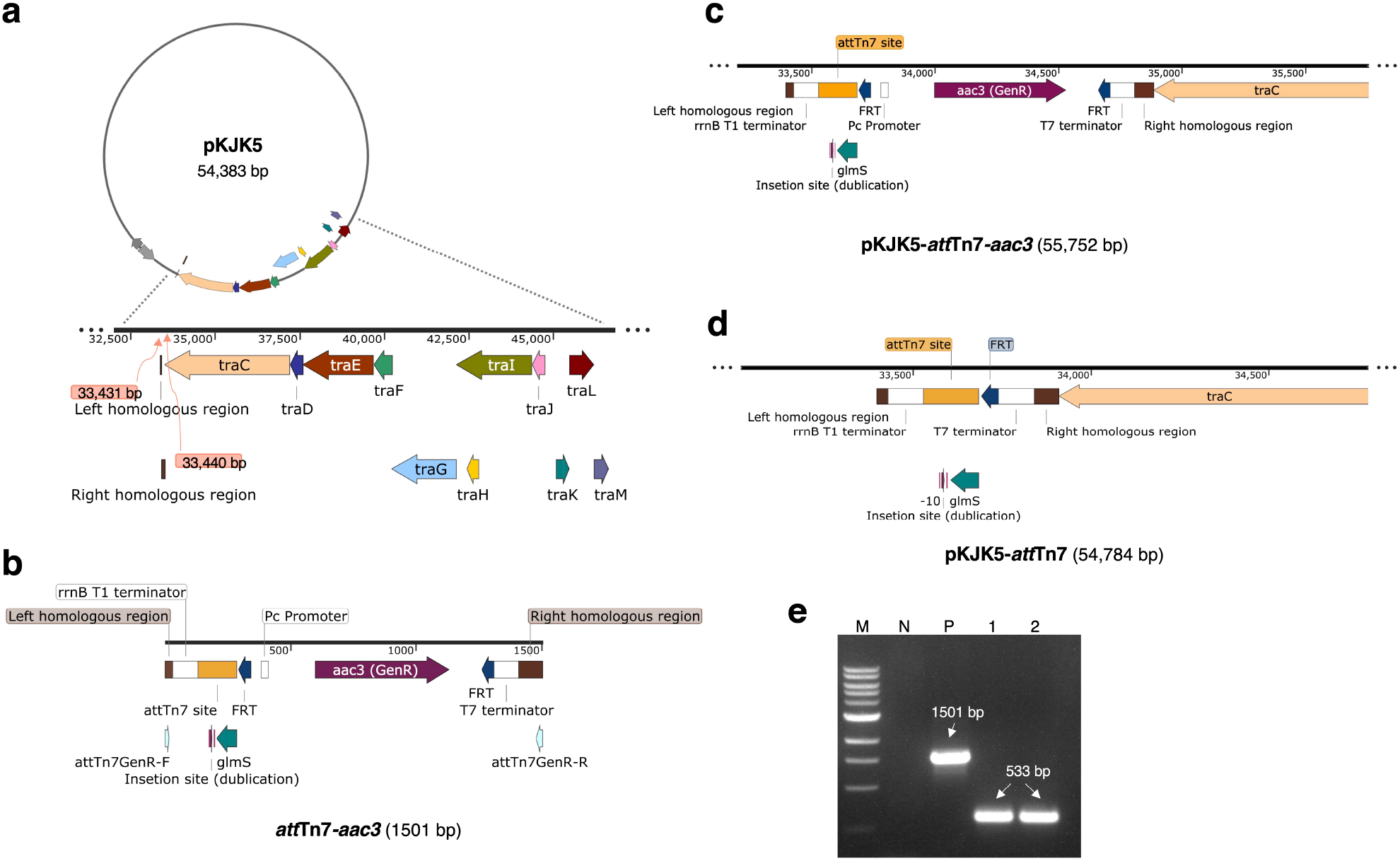
Stepwise engineering of pKJK5-*att*Tn7. a) The backbone of the conjugative broad-host-range IncP-1 plasmid pKJK5 and the region of insertion. b) Schematic diagram of the *att*Tn7 fragment used. c) Schematic diagram of the plasmid pKJK5-*att*Tn7*-aac3* after insertion of the *att*Tn7*-aac3* cassette into plasmid pKJK5. d) Schematic diagram of the plasmid pKJK5-*att*Tn7 after removal of resistance gene *aac3*. e) Colony PCR used to verify insertions. The primers *att*Tn7Gen^R^-F/R were used for PCR here. Lane M is DNA marker (NEB 1 Kb DNA Ladder), lane N shows the PCR product from negative control (ddH_2_O), lane P shows the PCR product from plasmid pKJK5-*att*Tn7-*aac3*, lanes 1 and 2 show the PCR products from two clones of pKJK5-*att*Tn7. Figures a), b), c) and d) were created with SnapGene software 6.0.2 (GSL Biotech).

### Construction of conjugation deficient plasmid pKJK5_CD_-*att*Tn7 and non-conjugative plasmid pKJK5_NC_-*att*Tn7

The λ Red homologous recombination method was also used to construct two additional versions of pKJK5-*att*Tn7. One that transfers at lower frequencies, referred to as pKJK5_CD_-*att*Tn7, and a non-conjugative version referred to as pKJK5_NC_-*att*Tn7. Plasmid pKD3 (Datsenko and Wanner, 2000) was used as template to amplified the chloramphenicol acetyltransferase gene (*cat*) including FRT-sites flanked by 40bp homology overhangs (1113 bp). The primers CD-*cat*-F/R and primers NC-*cat*-F/R were used here respectively. The *cat* fragment was integrated into the *traC* gene (pKJK5 nt position 36168-37226, AM261282) of pKJK5-*att*Tn7 using λ Red homologous recombination as described above. Hereafter, the plasmid pFLP2 was used to remove the *traC* region including the *cat* gene (pKJK5 nt position 33431-37226) hereby obtaining pKJK5_CD_-*att*Tn7. Similarly, in order to obtained pKJK5_NC_-*att*Tn7, the *cat* fragment was integrated upstream of the *traK* gene (pKJK5 nt position 43767-45046) of pKJK5-*att*Tn7, and then used the same method to remove the *traC-traJ* regions including the *cat* gene (pKJK5 nt position 33431-45046). Colony PCR with primer sets *att*Tn7Gen^R^-F and CD-*cat*-R/NC-*cat*-R were used to screen candidate colonies for pKJK5_CD_-*att*Tn7 and pKJK5_NC_-*att*Tn7, respectively. pKJK5_CD_-*att*Tn7 specific PCR products were purified and confirmed by Sanger sequencing (Eurofins Genomics). pKJK5_NC_-*att*Tn7 was whole-genome sequenced using the Illumina MiSeq platform. Sequencing libraries were prepared using the Nextera XT DNA Library Preparation Kit (Illumina, FC-131-1096) and sequenced with 2 × 250 base paired-end reads on the Illumina MiSeq platform (Illumina) according to the manufacturer’s protocol. Sequencing results were analyzed in CLC Genomic Workbench V7.5.1.

### Construction of mini-Tn7 pGRG36-based delivery vectors for *bla*-*sfGFP* complementation

Next, *sfGFP* and seven different β-lactamase genes were cloned into helper plasmid pGRG36 to subsequently be used as delivery into pKJK5-*att*Tn7 and pKJK5_NC_-*att*Tn7. *sfGFP* flanked by *tetA* was amplified by PCR from p-sfGFP, using primers psfGFP-F and psfGFP-R. This fragment was cloned into pGRG36 at the *Sma*I restriction site following the manufacturer’s protocol (*Sma*I, New England Biolabs and T4 DNA ligase, Thermo Scientific) resulting in pGRG36-sfGFP. Next, 7 β-lactamase genes originating from wildtype plasmids (*bla*_TEM-1_ (GeneBank Accession No. AY599232), *bla*_CTX-M-15_ (8ZF34, an wastewater isolate in our lab), *bla*_CMY-2_ (https://www.ncbi.nlm.nih.gov/biosample/?term=%20E.%20coli%2015083205), *bla*_NDM-5_ (an isolate from chicken cloaca), *bla*_KPC-2_ (ATCC^®^ BAA-1705™), *bla*_OXA-181_ (an isolate from hospital), *bla*_ampC_ (https://www.ncbi.nlm.nih.gov/biosample/?term=E.+coli+15093653)) were cloned into 7 different pGRG36*-sfGFP* vectors at the *Zra*I restriction site. Primers blaTEM1-F/R, blaCTXM15-F/R, blaCMY2-F/R, blaNDM5-F/R, blaKPC2-F/R, blaOXA181-F/R, blaampC-F/R were used to obtain genes *bla*_TEM-1_, *bla*_CTX-M-15_, *bla*_CMY-2_, *bla*_NDM-5_, *bla*_KPC-2_, *bla*_OXA-181_, *bla*_ampC_, respectively. Upon insertion of the *bla* genes the *tetA* gene was excised. Clones were selected on LB agar medium incubated at 30°C containing appropriate antibiotics. Verification was done by colony PCR using the aforementioned primer sets.

### Insertion of different *bla*-*sfGFP* cassettes into pKJK5-*att*Tn7 and pKJK5_NC_-*att*Tn7 using Tn7 transposition

First, the plasmids pKJK5-*att*Tn7 and pKJK5_NC_-*att*Tn7 were transformed by electroporation to *Escherichia coli* MG1655-*lacI*^*q*^*-mcherry* (Klümper et al., 2015) competent cells, where the chromosomal *att*Tn7-site is blocked by the *lacI*^*q*^*-mcherry* gene cassette, ensuring that Tn7 transposition inserts the *bla*-*sfGFP* cassettes into the *att*Tn7*-*site of pKJK5-*att*Tn7 and pKJK5_NC_-*att*Tn7.

Next, vectors pGRG36*-sfGFP* and pGRG36-*sfGFP-bla* (100-150 ng) were transformed by electroporation (1.8 kV for ∼5 ms) to *E. coli* MG1655-*lacI*^*q*^*-mcherry* competent cells containing the conjugative plasmid pKJK5-*att*Tn7 or pKJK5_NC_-*att*Tn7. Three colonies that grew on LB agar medium with Amp and Tet were selected for each of the 14 constructs. Next, these were grown in LB broth supplemented with 1 g/ml arabinose overnight at 30°C with shaking. Hereafter, cultures were re-streaked onto LB agar medium and incubated at 42°C overnight to cure cells for the different pGRG36 vectors. Next, clones were screened and verified by colony PCR with primers psfGFP-F/R and primers pGRG36QQ-F/R.

### Solid surface filter conjugation assay

Overnight cultures of donor strains and recipient strains were washed twice with PBS, and the OD_600_ of all cultures were adjusted to 0.5 with PBS. Next, donors and recipients were mixed at a ratio of 1:1 and immediately 50 μl was added onto a 0.2 μm mixed cellulose ester filter which was placed on top of LB agar medium. These were incubated for 20 hours at 37°C. After incubation, cells were transferred to a tube with 5 ml PBS by repeatedly pipetting cells off the filter. 100 μl of each of these suspensions were spread evenly onto agar plates with antibiotics as detailed below and incubated overnight at 37 °C. Transfer efficiencies were calculated as transconjugants per donor (Sørensen et al., 2005). LB agar plates containing Tet were used to count CFUs of the donors. LB agar plates containing Tet, Nal, and Rif were used to count CFUs of transconjugants. Three biological replicates were performed for each experiment and three technical replicates were made for each biological replicate.

### Growth curves

*E. coli* strains carrying plasmids pKJK5, pKJK5-*att*Tn7, pKJK5_CD_-*att*Tn7, or pKJK5_NC_-*att*Tn7 were cultured overnight in LB broth with Tet at 37°C. Then LB broth was used to make 10^5^-fold dilutions of the overnight cultures, and 200 μl diluted cultures were added to wells of a 96-well microtiter plate. Finally, the plate was incubated in a spectrophotometer (Bio Tek ELx808™ Absorbance Microplate Reader) at 37°C overnight, with continuously shaking, and OD_600_ was measured every 15 min. Nine replicates were done for each strain (three biological replicates each with three technical replicates).

### Flow cytometry

Flow cytometry was conducted to verify *sfGFP* expression using a BD FACS AriaIIIu (BD Biosciences) with a 488 nm excitation laser and FITC (530/30 nm band-pass filter) detector. Wild-type strain *E. coli MG1655* was used as a negative control. For sample treatment, dilute 5μl of the overnight cultured bacterial solution with 1ml of PBS. The threshold for forward scatter (FSC) was 1200, for side scatter (SSC) was 200, and the gating strategy was consistent with our previous study (Olesen et al., 2022). Data was acquired and analyzed using the BD FACSDiva software v.6.1.3.

### Antibiotic sensitivity

Inhibitory concentrations of antibiotics for all strains were determined using the agar dilution method (Wiegand et al., 2008). Overnight cultures were diluted to 10^4^ CFU/ml. 10 μl from each suspension were spotted onto LB agar plates containing a series of antibiotics (0-10 μg/ml MEM, 0-10 μg/ml CTX, 0-200 μg/ml AMP) to determine the minimum inhibitory concentration. The agar plates were incubated at 37 °C for 16 hours and the lowest antibiotic concentration that inhibited visible bacterial growth was regarded as inhibitory. These experiments were repeated independently three times.

### Statistical analysis

All the comparisons of mean values between conditions (*e.g*., OD_600_) were performed using two-sided Student’s t-tests calculated using R 4.0 (Team, 2013). Growth curves were analyzed with the R package Growthcurver (Sprouffske and Wagner, 2016).

## Results

### Plasmid pKJK5-*att*Tn7

Here, an *att*Tn7-site was introduced into conjugative IncP plasmid pKJK5 (**Fig. 1a**). Based on the genome sequence of pKJK5, we chose to insert the *att*Tn7-site between nt 33431-33440 (**Fig. 1a**), as this was at the interface between plasmid backbone and accessory genes with little risk of functional disruption. The region may have encoded a terminator associated with *traC*, however, a new terminator was added with the *att*Tn7 fragment (**Fig. 1b**). The inserted *att*Tn7*-aac3* fragment was flanked on both side with strong terminators to diminish expression spillover both from flanking genes and from genes to be inserted into the *att*Tn7-site (**Fig. 1c**). *aac3* was introduced alongside the *att*Tn7-site and was subsequentially removed by Flp recombinase-mediated excision via *FRT*-sites that flanked *aac3* (**Fig. 1d** and **Fig. 2a**). Colony PCR and full genome sequencing verified pKJK5-*att*Tn7 had been constructed successfully (**Fig. 1e**).

**Fig. 2.**
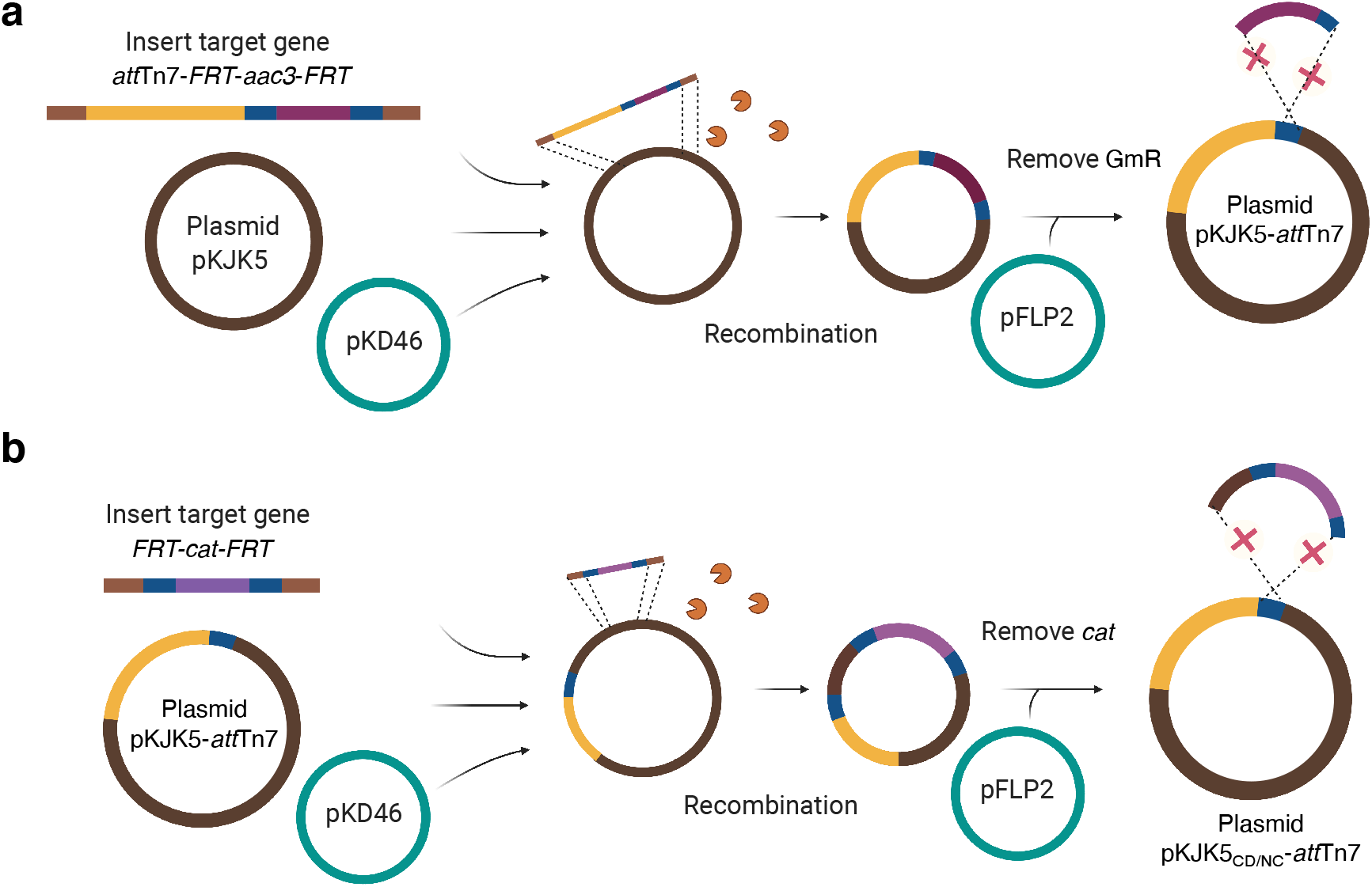
Schematic diagram of the construction of plasmids pKJK5-*att*Tn7, pKJK5_CD_-*att*Tn7 and pKJK5_NC_-*att*Tn7. λ Red homologous recombination was used when constructing plasmid pKJK5-*att*Tn7. The helper plasmid pKD46 encoding the Red homologous recombinases was used to integrate the PCR product *att*Tn7*-FRT*-*aac3*-*FRT* into pKJK5. The fragment *att*Tn7*-FRT*-*aac3*-*FRT* contained the gentamicin resistance gene (*aac3*) and facilitated the selection of positive clones. Vector pFLP2 which facilitates Flp recombination of FRT-sites eliminated the gentamicin resistance gene. λ Red homologous recombination was used when constructing plasmid pKJK5_CD_-*att*Tn7 and pKJK5_NC_-*att*Tn7. Insertion of the fragment *FRT*-*cat*-*FRT* into the plasmid pKJK5-*att*Tn7 was performed in a similar way as a). With the help of the *FRT*-site left in pKJK5-*att*Tn7, Flp recombination removed *traC-cat* from pKJK5_CD_-*att*Tn7 and the *traC*-*traJ-cat* region from pKJK5_NC_-*att*Tn7. Figures were created with BioRender.com.

### Conjugation deficient plasmid pKJK5_CD_-*att*Tn7 and non-conjugative plasmid pKJK5_NC_-*att*Tn7

Next, we removed *traC* and the *traC-traJ* region of pKJK5-*att*Tn7 to create a version that conjugates at lower frequencies (pKJK5_CD_-*att*Tn7) and a non-conjugative version (pKJK5_NC_-*att*Tn7), respectively (Fig. 3a & b). pKJK5_CD_-*att*Tn7 and pKJK5_NC_-*att*Tn7 can be used as comparative controls in various experimental setups. *FRT*-*cat*-*FRT* was introduced upstream of *traC* for pKJK5_CD_-*att*Tn7 (**Fig. 3a**) and upstream *traI* (and *traK*) (**Fig. 3b**) using λ Red recombination. Hereafter, Flp recombination was used to removed regions between the *FRT*-site associated with *att*Tn7 and those introduced with *cat*. This resulted in the deletion of the *traC-FRT-cat-FRT* region constructing pKJK5_CD_-*att*Tn7 and the deletion of *traCDEFGHI-FRT-cat-FRT* region constructing pKJK5_NC_-*att*Tn7 (**Fig. 2b**). Removal of *traC* in pKJK5_CD_-*att*Tn7 and *traC-I* in pKJK5_NC_-*att*Tn7 was verified by colony PCRs (**Fig. 3d**).

**Fig. 3.**
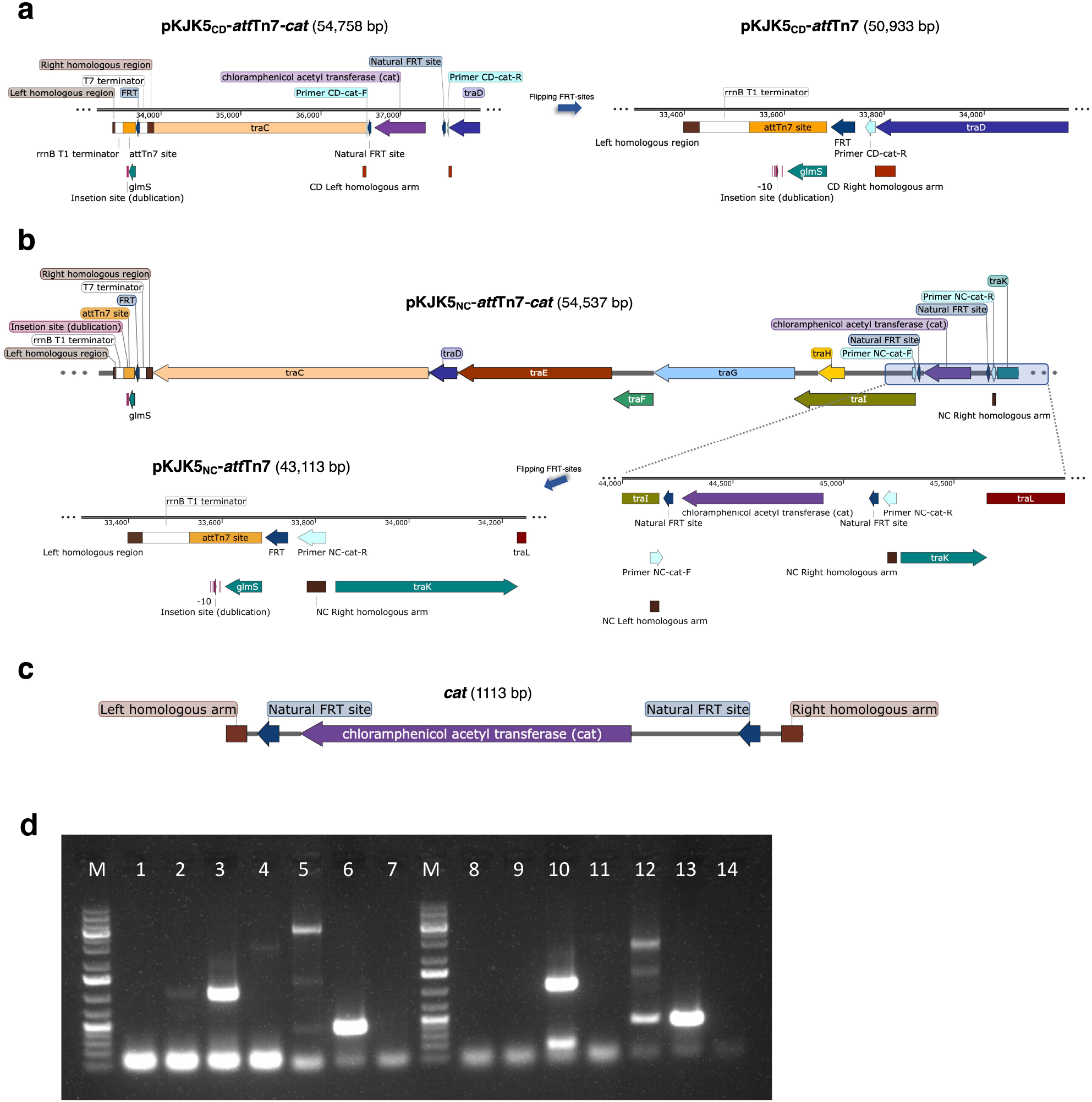
Stepwise engineering of plasmids pKJK5_CD_-*att*Tn7 and pKJK5_NC_-*att*Tn7. a) *FRT*-*cat*-*FRT* was inserted into pKJK5*-att*Tn7 upstream *traC*. Hereafter, Fpl recombination was used to remove *traC-FRT-cat-FRT* constructing conjugation deficient plasmid pKJK5_CD_*-att*Tn7 b) *FRT*-*cat*-*FRT* was inserted into pKJK5*-att*Tn7 upstream *traI*. Fpl recombination was used to remove *traCDEFGHI-FRT-cat-FRT* constructing non-conjugative plasmid pKJK5_NC_*-att*Tn7. Schematic diagram of the *FRT*-*cat*-*FRT* fragment. d) Colony PCR to verify the construction of pKJK5_CD_*-att*Tn7 and pKJK5_NC_*-att*Tn7. Lanes M are 1 kb DNA ladders (Thermo Scientific™ GeneRuler™ 1 kb Plus DNA Ladder). Lanes 1, 7, 8, and 14 show the PCR product from negative control (ddH_2_O). Lanes 1, 2, 3, and 4 show the PCR product from pKJK5-*att*Tn7, pKJK5_CD_-*att*Tn7*-cat*, pKJK5_CD_-*att*Tn7, and negative control (ddH_2_O) with primers CD-*cat*-F/R, respectively. Lanes 5, 6, and 7 show the PCR product from plasmid pKJK5_CD_-*att*Tn7*-cat*, pKJK5_CD_-*att*Tn7, and negative control (ddH_2_O) with primers *att*Tn7Gen^R^-F and CD-*cat*-R, respectively. Lanes 8, 9, 10, and 11 show the PCR product from plasmid pKJK5-*att*Tn7, pKJK5_NC_-*att*Tn7*-cat*, pKJK5_NC_-*att*Tn7, and negative control (ddH_2_O) with primers NC-*cat*-F/R, respectively. Lanes 12, 13, and 14 show the PCR product from plasmid pKJK5_NC_-*att*Tn7*-cat*, pKJK5_NC_-*att*Tn7, and negative control (ddH_2_O) with primers *att*Tn7Gen^R^-F and NC-*cat*-R, respectively. Figures a), b) and c) were created with SnapGene software 6.0.2 (GSL Biotech).

### Growth and conjugal transfer frequencies

Growth curves were made of strains *E. coli* MG1655//pKJK5, *E. coli* MG1655//pKJK5-*att*Tn7, *E. coli* MG1655//pKJK5_CD_-*att*Tn7, and *E. coli* MG1655//pKJK5_NC_-*att*Tn7 (**Fig. 4a**). No significant difference was found between growth rates or the doubling times between the strains (*P* < 0.05, one-way ANOVA with post-hoc Tukey test).

**Fig. 4.**
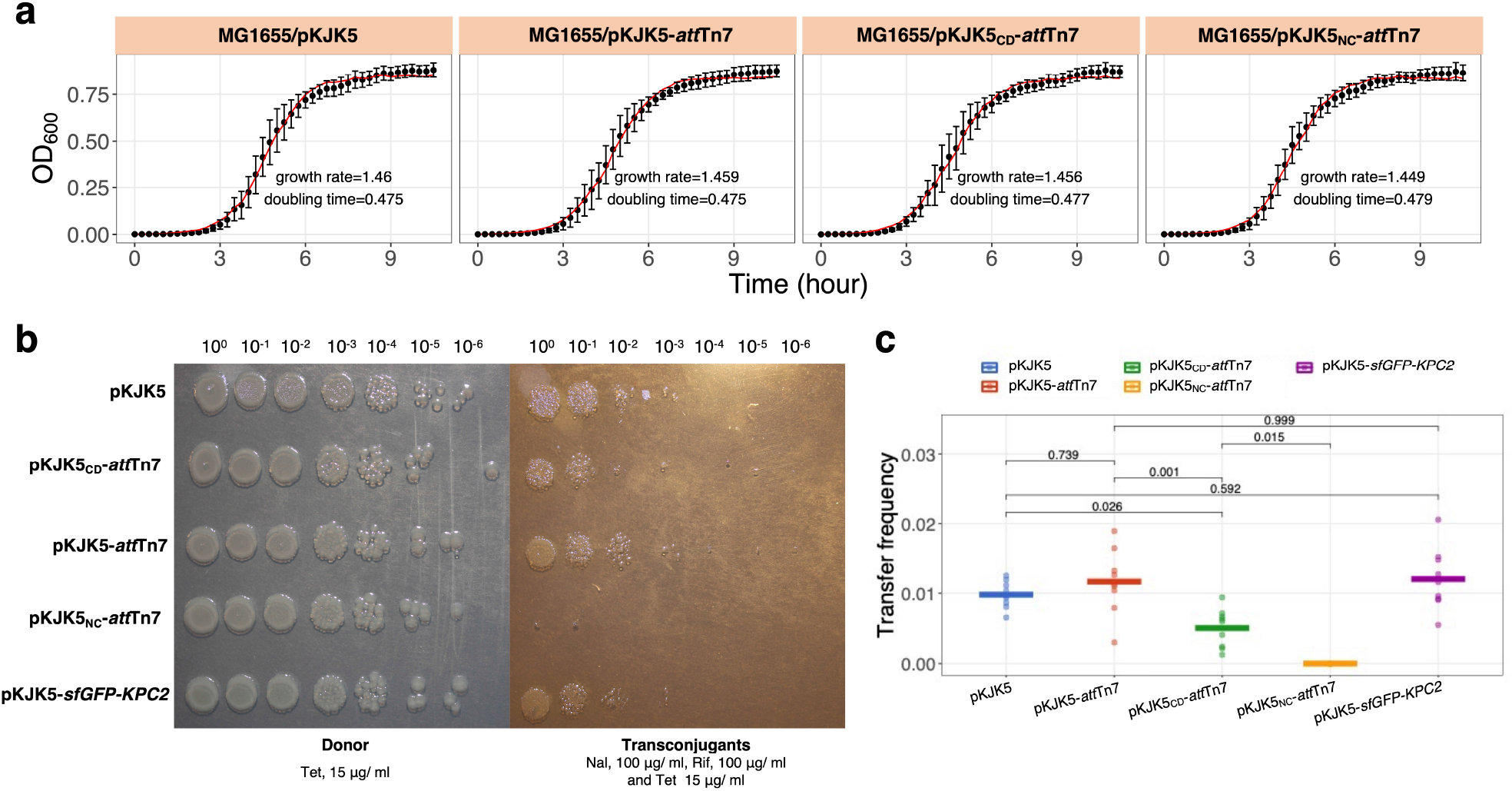
Growth and conjugal transfer frequencies. a) Growth curves of *E. coli* MG1655 strains carrying pKJK5 or one of the 3 derived plasmids in LB broth (n =9). The black dots represent the mean, the black line the standard deviation, and the red line the fitted growth curve. b) CFUs showing the transfer efficiency of the 5 plasmids. Representative images that show similar numbers of donors and transconjugants in experiments with pKJK5, pKJK5-*att*Tn7 and pKJK5-*att*Tn7::*bla*_*KPC-2*_-*sfGFP*. The number of transconjugants was lower for pKJK5_CD_-*att*Tn7 and no conjugational transfer occurred of pKJK5_NC_-*att*Tn7, even though the number of donors were similar. c) Transfer frequencies (Tranconjugants/Donors) of the 5 different plasmids. Dots are individual replicates (n = 9) and lines are the mean. *P-* values were derived from a one-way ANOVA followed with post-hoc Tukey test.

Next, we used the strains *E. coli* MG1655//pKJK5, *E. coli* MG1655//pKJK5-*att*Tn7, *E. coli* MG1655//pKJK5_CD_-*att*Tn7, and *E. coli* MG1655//pKJK5_NC_-*att*Tn7 as donors, and strain *E. coli* MG1655-Kan^R^-Rif^R^-Nal^R^ as recipient, to measure conjugation transfer frequencies (**Fig**.**4b-c**). We found that the insertion of the *att*Tn7-site did not affect conjugal transfer as transfer frequencies of wildtype plasmid pKJK5 and pKJK5-*att*Tn7 were similar (*P* = 0.739, one-way ANOVA with post-hoc Tukey test). The transfer frequency of *traC* deletion plasmid pKJK5_CD_-*att*Tn7 was significantly reduced (*P* < 0.05, one-way ANOVA with post-hoc Tukey test) rendering it conjugation deficient. No transconjugants were observed in transfer experiments with pKJK5_NC_-*att*Tn7 showing that this plasmid had lost its ability to conjugate.

### Gene complementation at the *att*Tn7-site of the constructed plasmids

To test the efficacy of the *att*Tn7 complemented plasmids, 7 different β-lactamase genes (*bla*_TEM-1_, *bla*_CTX-M-15_, *bla*_CMY-2_, *bla*_NDM-5_, *bla*_KPC-2_, *bla*_OXA-181_, *bla*_ampC_) flanked by *sfGFP* were inserted into pKJK5-*att*Tn7 using the mini-Tn7 delivery vector PGRG36. Since the *att*Tn7 site typically also exists in the bacterial chromosome, *E. coli* MG1655-*lacI*^*q*^*-mcherry* (Klümper et al., 2015) was used as background when performing complementation. In *E. coli* MG1655-*lacI*^*q*^*-mcherry* the chromosomal *att*Tn7*-*site was occupied by the *lacI*^*q*^*-mcherry* cassette preventing further chromosomal transposition events. PCR, gel electrophoresis, and Sanger sequencing was used to verify that the plasmids has been successfully constructed (**Fig. 5a**). The different β-lactamase genes and *sfGFP* were also inserted into the non-conjugative plasmid pKJK5_NC_-*att*Tn7, illustrating the high efficacy of gene complementation into the *att*Tn7*-*sites located on the constructed plasmids presented here (**Fig. 5a**). Next, to verify the expression of the inserted genes, inhibitory concentrations of the different strains were tested (**Table 3**). The inhibitory concentrations towards relevant β-lactamases of all strains carrying the different *bla* genes in pKJK5-*att*Tn7, were significantly higher than those of the wildtype *E. coli* MG1655 (**Table 3**). To test the expression of *sfGFP*, green fluorescence was measured by flow cytometry using *E. coli* MG1655-*lacI*^*q*^*-mcherry//*pKJK5-*att*Tn7::*sfGFP-KPC2* as a representative (**Fig. 5b-c**). Here the constitutively expressed *lacI*^*q*^ represses the P_*A1-O4/O3*_ promoter of *sfGFP*. To counteract this 1 mM Isopropyl β-D-1-thiogalactopyranoside (IPTG) was added to relieve the repression of *sfGFP* by *lacI*^*q*^ (Lanzer and Bujard, 1988; Dahlberg et al., 1998). Lastly, we tested if the *bla*_*KPC-2*_-*sfGFP* insertion influenced conjugation frequencies of plasmid pKJK5 (**Fig. 4b-c**). We found that the *bla*_*KPC-2*_ and *sfGFP* were expressed as expected and transfer frequencies were similar to the wildtype plasmid pKJK5 (*P* = 0.592, one-way ANOVA with post-hoc Tukey test).

**Table 3.**
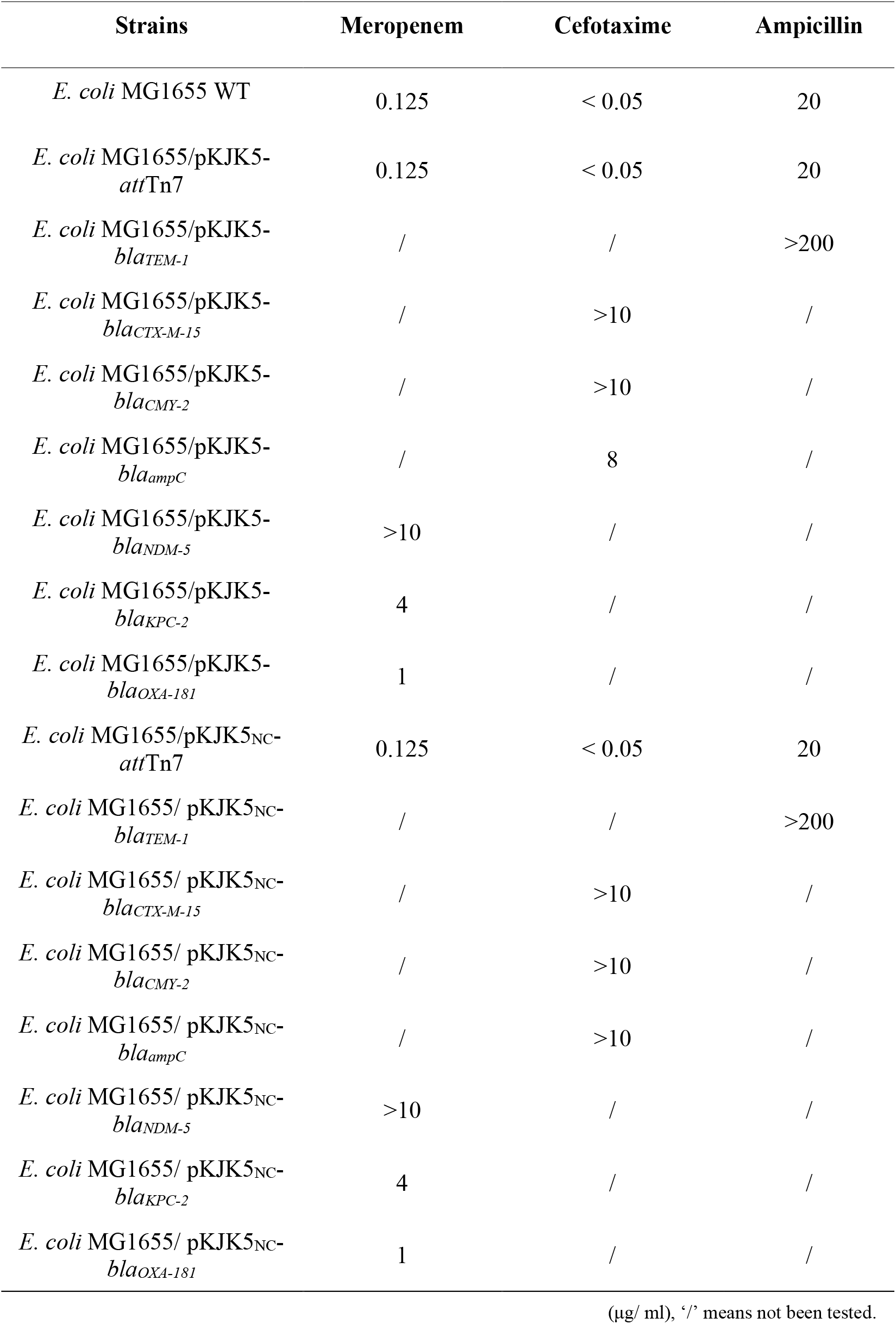
The minimum inhibitory concentration (MIC) of strains.

| Strains | Meropenem | Cefotaxime | Ampicillin |
| --- | --- | --- | --- |
| *E. coli* MG1655 WT | 0.125 | < 0.05 | 20 |
| *E. coli* MG1655/pKJK5-<br>*attTn7* | 0.125 | < 0.05 | 20 |
| *E. coli* MG1655/pKJK5-<br>*bla*<sub>TEM-1</sub> | / | / | >200 |
| *E. coli* MG1655/pKJK5-<br>*bla*<sub>CTX-M-15</sub> | / | >10 | / |
| *E. coli* MG1655/pKJK5-<br>*bla*<sub>CMY-2</sub> | / | >10 | / |
| *E. coli* MG1655/pKJK5-<br>*bla*<sub>ampC</sub> | / | 8 | / |
| *E. coli* MG1655/pKJK5-<br>*bla*<sub>NDM-5</sub> | >10 | / | / |
| *E. coli* MG1655/pKJK5-<br>*bla*<sub>KPC-2</sub> | 4 | / | / |
| *E. coli* MG1655/pKJK5-<br>*bla*<sub>OXA-181</sub> | 1 | / | / |
| *E. coli* MG1655/pKJK5<sub>NC</sub>-<br>*attTn7* | 0.125 | < 0.05 | 20 |
| *E. coli* MG1655/ pKJK5<sub>NC</sub>-<br>*bla*<sub>TEM-1</sub> | / | / | >200 |
| *E. coli* MG1655/ pKJK5<sub>NC</sub>-<br>*bla*<sub>CTX-M-15</sub> | / | >10 | / |
| *E. coli* MG1655/ pKJK5<sub>NC</sub>-<br>*bla*<sub>CMY-2</sub> | / | >10 | / |
| *E. coli* MG1655/ pKJK5<sub>NC</sub>-<br>*bla*<sub>ampC</sub> | / | >10 | / |
| *E. coli* MG1655/ pKJK5<sub>NC</sub>-<br>*bla*<sub>NDM-5</sub> | >10 | / | / |
| *E. coli* MG1655/ pKJK5<sub>NC</sub>-<br>*bla*<sub>KPC-2</sub> | 4 | / | / |
| *E. coli* MG1655/ pKJK5<sub>NC</sub>-<br>*bla*<sub>OXA-181</sub> | 1 | / | / |

(µg/ ml), ‘/’ means not been tested.

**Fig. 5.**
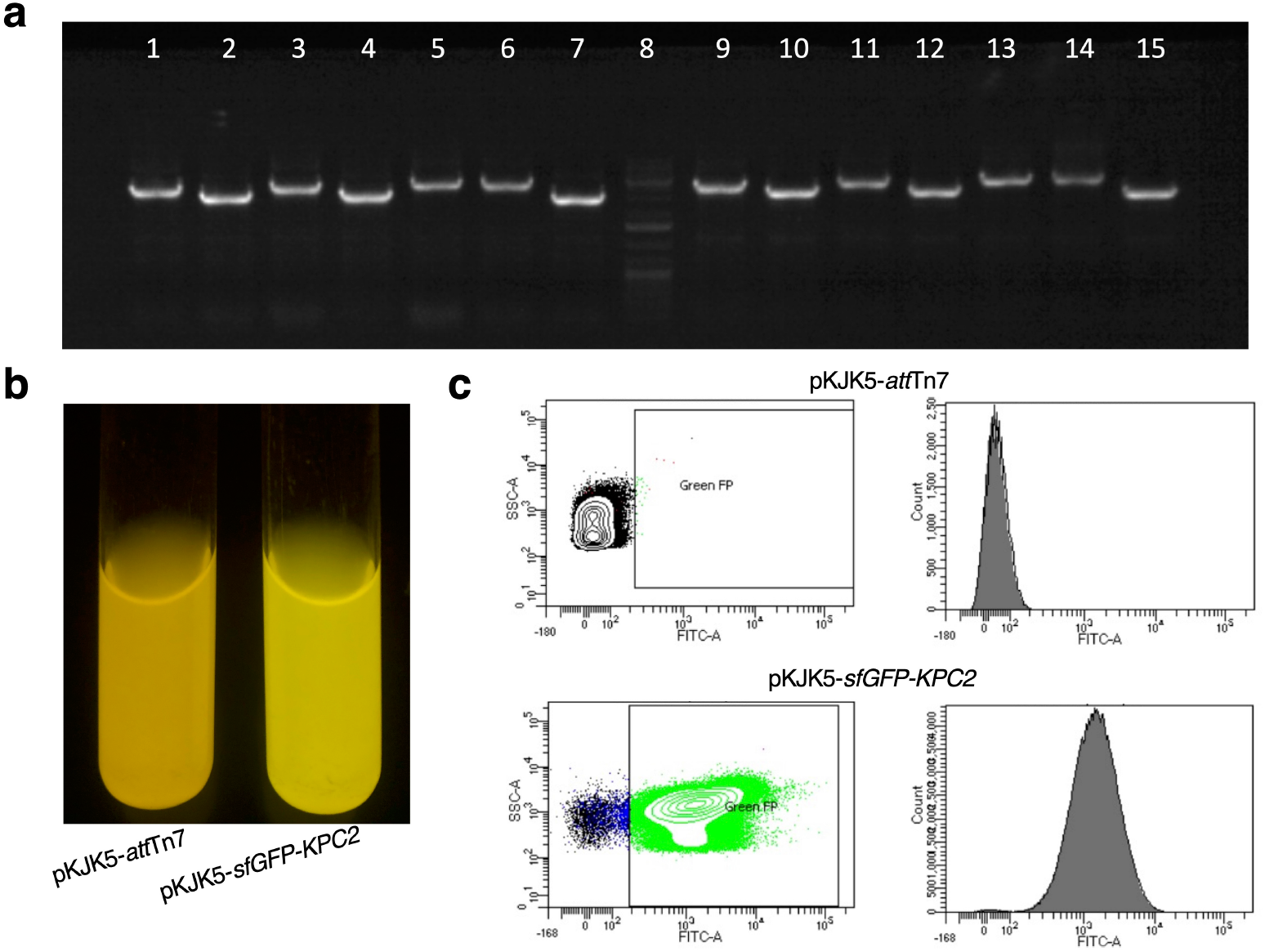
Verification of insertion of *bla-sfGFP* cassettes into pKJK5-*att*Tn7 and pKJK5_NC_-*att*Tn7. a) PCR verification of the correct insertion of the different *bla-sfGFP* cassettes into pKJK5-*att*Tn7 and pKJK5_NC_-*att*Tn7. Lanes 1, 2, 3, 4, 5, 6, and 7 verifies insertion of *bla*_*KPC-2*_, *bla*_*NDM-5*_, *bla*_*OXA-181*_, *bla*_*CTXM-15*_, *bla*_*ampC*_, *bla*_*CMY-2*_, and *bla*_*TEM-1*_, respectively, into pKJK5-*att*Tn7. Lane 8 a 1 kb DNA Ladder (Thermo Scientific™ GeneRuler™ 1 kb Plus DNA Ladder). Lanes 9, 10, 11, 12, 13, 14, and 15 verifies insertion of *bla*_*KPC-2*_, *bla*_*NDM-5*_, *bla*_*OXA-181*_, *bla*_*CTXM-15*_, *bla*_*ampC*_, *bla*_*CMY-2*_, and *bla*_*TEM-1*_, respectively, into pKJK5_NC_-*att*Tn7. b) Image showing green fluorescence under ultraviolet light due to expression of the *sfGFP* by *E. coli* MG1655//pKJK5-*att*Tn7::*bla*_*KPC-2*_-*sfGFP. E. coli* MG1655//pKJK5-*att*Tn7 was used as a negative control. c) Green fluorescence due to expression of the *sfGFP* by *E. coli* MG1655//pKJK5-*att*Tn7::*bla*_*KPC-2*_-*sfGFP* detected by flow cytometry. *E. coli* MG1655//pKJK5-*att*Tn7 was used as a negative control.

## Discussion

Here we present a suite of engineered plasmids that were developed to advance research on HGT and plasmid biology. We introduced an *att*Tn7-site into conjugative broad-host-range IncP-1 plasmid pKJK5 using the λ phage derived Red recombination system. Hereafter, we demonstrate the ease at which genes of interest can be inserted into the *att*Tn7-site with high efficacy using the bacterial mini-Tn7 transposon site-specific recombination system. We utilized vector pGRG36 which contains a multiple cloning site for easy complementation, and it allows insertion of large fragments. It is known that Tn7 possesses transposition immunity, which counteracts Tn7 from transposing into *att*Tn7 sites already occupied by Tn7 (DeBoy and Craig, 1996), reducing the chance of at least 10^4^ fold (McKenzie and Craig, 2006). The construction of the 14 different plasmids with different *bla* genes and *sfGFP* (pKJK5-*sfGFP*^-^*bla)* illustrates that, e.g., when wanting to study a series of functional genes in the context of a conjugative plasmid, the introduction of an *att*Tn7-site reduced time spend engineering and overall costs.

The mini-Tn7-based transposon system is typically used for the insertion of single gene cassettes, however, innovative approached such as high-throughput chromosomal-barcoding to track the evolutionary dynamics of *E. coli* subpopulations has recently been implemented (Jasinska et al., 2020) and illustrated the broad potential of these systems. The λ Red recombination system can also be used on its own to engineer wildtype plasmids (Anjum et al., 2018), however, the recombination efficiency is often limited by the size of the DNA insert (Doron et al., 2018). Besides, our experience is that using the λ Red recombination system can be quite challenging when attempting to engineer wildtype plasmids despite extended experience with these systems. Novel systems like INTEGRATE (Vo et al., 2021) may prove well suited for engineering wildtype plasmids.

The approach presented here can be seen as a proof-of-concept as any *att*-site and associated transposons/integrases could be used to generate similar systems, not only based on conjugative wildtype plasmids but any mobile genetic element. For some applications one might want to use less widespread *att*-site/integration systems (*e.g*., the CTX system), modular *att*-sites/integration systems (*e.g*., gateway systems), or others (Rajeev et al., 2007; Merrick et al., 2018). Here the *att*Tn7-site and the mini-Tn7 system were chosen partially because it, in addition to the pKJK5 derived plasmids described here, also enables us to compliment chromosomal *att*Tn7-sites (Choi et al., 2005), which can be valuable for comparative purposes. For the same reason, we constructed the conjugation deficient plasmid pKJK5_CD_-*att*Tn7 and non-conjugative plasmid pKJK5_NC_-*att*Tn7.

## Data availability

Source data are provided with this paper.

## Acknowledgements

We thank Valeria Bortolaia at the Technical University of Denmark for providing strains *E. coli* 15083205 and *E. coli* 15093653 used as templates for amplification of genes *bla*_*CMY-2*_ and *bla*_*ampC*_, respectively. We thank Qiue Yang and Timothy R. Walsh team from Oxford university for providing the genes *bla*_*NDM-5*_ and *bla*_*OXA-181*_. We thank Ana Filipa Silva for kindly providing the genes *bla*_*KPC-2*_. We would like to thank the Lundbeck Foundation (JSM, R250-2017-1392) and Villum Fonden (JSM, 00028304) for supporting this study and China Scholarship Council for funding QW.

## Author contributions

QW, JSM and SJS designed the study. QW and JSM carried out the study. JSM, SJS, and QW contributed to the concept and interpretation of the data. QW drafted the manuscript. AKO contributed to the manuscript preparation. LM contributed to the plasmid sequencing.

